# Inter- and intra-domain functional redundancy in the rumen microbiome during plant biomass degradation

**DOI:** 10.1101/254672

**Authors:** Andrea Söllinger, Alexander Tøsdal Tveit, Morten Poulsen, Samantha Joan Noel, Mia Bengtsson, Jörg Bernhardt, Anne Louise Frydendahl Hellwing, Peter Lund, Katharina Riedel, Christa Schleper, Ole Højberg, Tim Urich

**Affiliations:** Department of Ecogenomics and Systems Biology, University of Vienna, Althanstraße 14, 1090 Vienna, Austria; Institute of Microbiology, University of Greifswald, Felix-Hausdorff-Straße 8, 17487 Greifswald, Germany; Department of Arctic and Marine Biology, UiT, The Arctic University of Norway, Hansine Hansens veg 18, 9019 Tromsø, Norway; Department of Animal Science, Aarhus University, Blichers Alle 20, 8830 Tjele, Denmark

**Keywords:** metatranscriptomics, methane, rumen, microbiome, carbohydrate active enzymes, volatile fatty acids, methanogenesis, archaea, *Methanomassiliicoccales*

## Abstract

**Background:** Ruminant livestock is a major source of the potent greenhouse gas methane (CH_4_), produced by the complex rumen microbiome. Using an integrated approach, combining quantitative metatranscriptomics with gas- and volatile fatty acid (VFA) profiling, we gained fundamental insights into temporal dynamics of the cow rumen microbiome during feed degradation.

**Results:** The microbiome composition was highly individual and remarkably stable within each cow, despite similar gas emission and VFA profiles between cows. Gene expression profiles revealed a fast microbial growth response to feeding, reflected by drastic increases in microbial biomass, CH_4_ emissions and VFA concentrations. Microbiome individuality was accompanied by high inter- and intra-domain functional redundancy among pro- and eukaryotic microbiome members in the key steps of anaerobic feed degradation. Methyl-reducing but not CO_2_-reducing methanogens were correlated with increased CH_4_ emissions during plant biomass degradation.

**Conclusions:** The major response of the rumen microbiome to feed intake was a general growth of the whole community. The high functional redundancy of the cow-individual microbiomes was possibly linked to the robust performance of the anaerobic degradation process. Furthermore, the strong response of methylotrophic methanogens is suggesting that they might play a more important role in ruminant CH_4_ emissions than previously assumed, making them potential targets for CH_4_ mitigation strategies.

## Background

Ruminant animals are the dominant large herbivores on Earth. Their evolutionary success is partly due to their tight symbiotic associations with commensal microorganisms (their microbiome) that enables them to utilise otherwise indigestible plant biomass as food sources [1]. Since their domestication in the Holocene, ruminants, in particular cows, have provided humankind with various important goods. However, agricultural farming of cows is also a major source of the potent greenhouse gas (GHG) methane (CH_4_), having a global warming potential 28 times higher than carbon dioxide [2].

Cows possess a complex digestive system including a four-compartment stomach, with the largest compartment being the rumen [3], a big anaerobic fermentation chamber harbouring the complex microbiome responsible for the anaerobic degradation of ingested plant biomass. During microbial hydrolysis and fermentation of plant fibres volatile fatty acids (VFA) are produced, which serve as the main energy source of the animal [4]. A prominent end-product of microbial degradation is CH_4_, produced by methanogenic Archaea. Individual cows, respectively their symbiotic methanogens, produce up to 500 L of CH_4_ per day [5], making ruminant livestock one major anthropogenic CH_4_ source [6]. Due to an increasing human world population, milk and meat demands are expected to double by 2050 [7], making the development of sustainable and productive animal farming systems a major challenge in agriculture [8]. CH_4_ mitigation strategies are not only of ecological, but also of economic importance as ruminant CH_4_ emissions represent an energy loss of 2 - 12 % for the animal [5, 8].

Since the times of the pioneering work of Hungate and others [9, 10, 11, 12], microbiologists have made large efforts to understand the structure-function relationships in the complex rumen microbiome, identifying the microorganisms that participate in certain steps of the anaerobic degradation pathway. More recently, the application of cultivation-independent molecular techniques has helped to uncover the high diversity of bacteria, archaea and eukaryotes residing in the rumen and factors affecting community composition (e.g. [13]). In addition, the usage of meta-omics techniques has paved the way for a better understanding of the rumen ecosystem and the microbial metabolic potential and activity in the rumen (reviewed by [14]). These studies have revealed differences in rumen microbiome structure between low and high CH_4_ emitting cows (e.g. [15, 16]) and the effects of different diets on ruminant CH_4_ emissions (e.g. [17, 18, 19]). New insights were also gained by identification of new members of functional groups e.g. new fibrolytic and methanogenic community members [20, 21, 22, 23, 24]. Furthermore, the importance of diurnal microbiome dynamics for the understanding of VFA, H_2_ and CH_4_ production in the rumen was pointed out recently [25].

Despite these major advances, a holistic understanding of the rumen microbiome is still lacking, including answers to rather simple questions such as “who is doing what and when during feed degradation?”. Such a fundamental understanding of the rumen ecosystem, as it was proposed by Hungate already in the early 1960s [11], can help to specifically manipulate the rumen microbiome, to lower CH_4_ emissions, without hampering animal productivity, milk and meat quality or being harmful to the animal [14, 26].

To obtain a more comprehensive view on the activity of the rumen microbiome during plant biomass degradation, we performed a longitudinal metatranscriptomics study of microbiome dynamics in lactating cows. We aimed at identifying the active pro- and eukaryotic microbiome members and define their function in the key steps of anaerobic polysaccharide degradation and CH_4_ production. We hypothesized that the microbiome exhibits a defined successional pattern, reflecting a cascade of hydrolytic, fermentative and methanogenic steps, accompanied by distinct VFA and gas emission patterns. Based on a previous metatranscriptomic study from our lab [24] and work of others ([27] and references therein), we hypothesized that the recently discovered *Methanomassiliicoccales* are substantial contributors to ruminant CH_4_ emissions and will therefore show high activity after ruminant feed intake.

Quantitative metatranscriptomics combined with gas- and VFA profiling enabled linking rumen microorganisms and their transcript profiles to processes. We show extensive inter- and intra-domain functional redundancy among microorganisms at several steps of the anaerobic degradation pathway.

## Results

### Temporal dynamics of feed digestion

To investigate the effect of feed intake on CH_4_ production by the rumen microbiome we conducted a diurnal feeding experiment over four days and measured CH_4_, CO2 and H_2_ emissions of four individual lactating Holstein cows on day four in open circuit respiration chambers (Fig. 1, Supplementary Table S1 and S2). Immediately after the morning feeding, CH_4_ and CO2 emissions approximately doubled (23.4 ± 2.2 L h^-1^ CH_4_ and 296.3 ± 11.0 L CO2 h^-1^) with all animals showing similar dynamics and magnitude of gas production (Fig. 1b). The emissions dropped to before-feeding levels four to six hours after feed intake. H_2_ was only detectable during the first hour after feeding started (Fig. 1b) indicative of highly active H_2_-producing primary and secondary fermenters providing excessive substrate for hydrogenotrophic methanogens. Similar dynamics in gas emissions were observed during afternoon feeding (Supplementary Figure SI). Likewise, the concentration of VFA in rumen fluid samples increased with peak concentrations measured three and two hours after start of the morning and the afternoon feeding, respectively (Fig. 1c), similar to [25]. However, compared to the gas emission profiles VFA pools were more variable in terms of magnitude and temporal dynamics between the four cows. The immediate accumulation of the fermentation products H_2_, CO_2_ (i.e. substrates for methanogenesis) and VFA after feeding indicated a fast physiological response of the rumen microbiome to feed intake, with enhanced fermentation rates leading to increased methanogenesis rates. Furthermore, a transient increase of RNA in the rumen fluid was observed, which we consider a proxy for active microbiome biomass. 34.1 ± 6.5, 69.2 ± 10.3, 70.0 ± 13.9 and 37.0 ± 6.3 μg RNA were extracted per gram rumen fluid at tO, tl, t3 and t5, respectively (Fig. 1d). Rumen fluids for VFA quantification and RNA extraction were sampled prior to gas measurements, as it was not possible to sample during respiration chamber measurements. The similar patterns in gases, VFA and RNA profiles reflected a similar behaviour of the cows during the animal feeding trial (Supplementary Table S2). Taken together, GHG emissions, VFA production and RNA content indicated a consistent and fast growth response of the rumen microbiome and strong temporal dynamics on process level (Fig. 1b-d, Supplementary Figure S2) within each individual cow.

**Figure 1.**
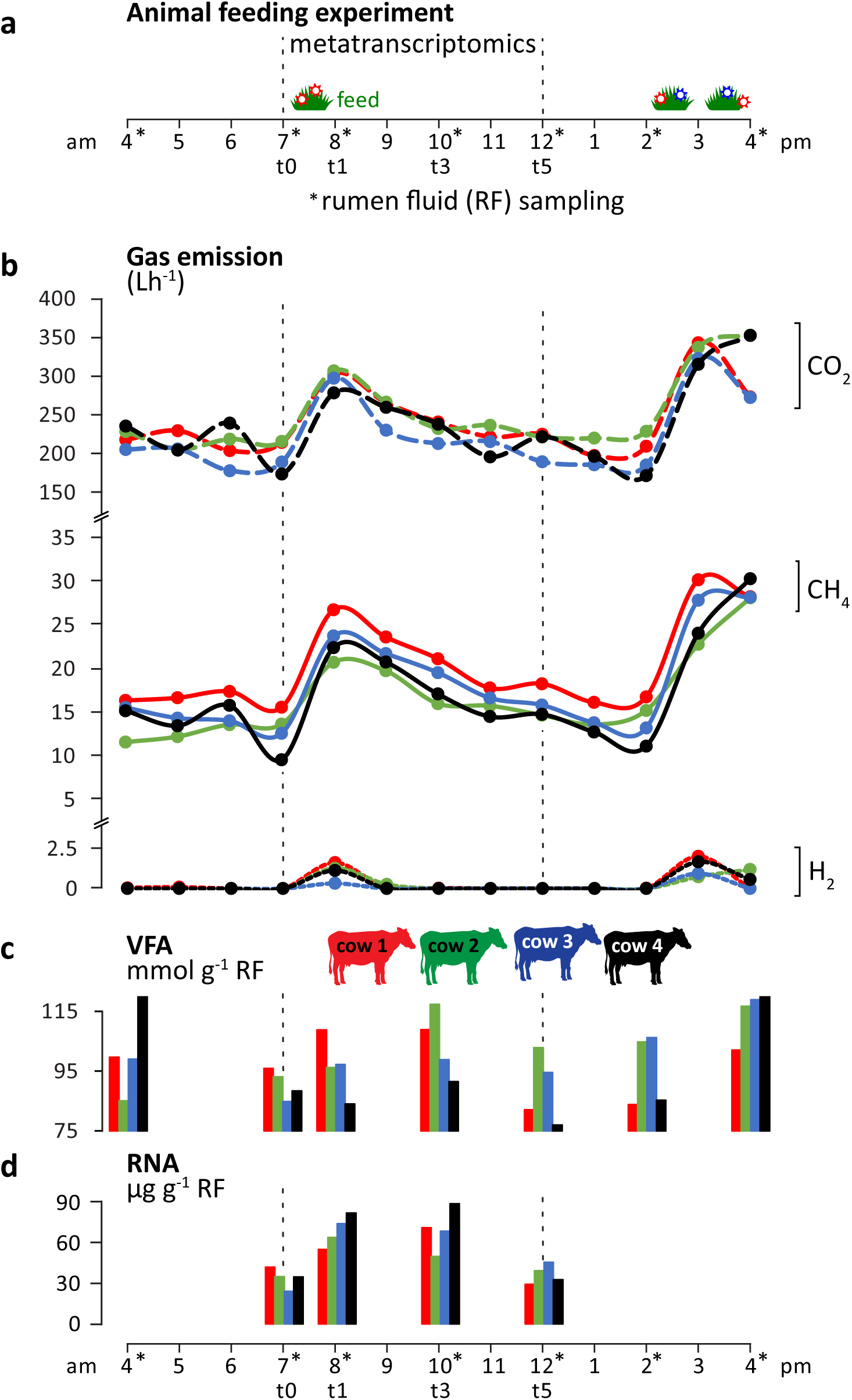
Ruminant gas emissions and volatile fatty acid (VFA) production. (a) Overview of the animal feeding trial during twelve hours (4 am - 4 pm) of sampling; for more details see Methods section, (b) Carbon dioxide (CO_2_), methane (CH_4_) and hydrogen (H_2_) emissions measured using open-circuit respiration chambers, (c), (d) Total VFA concentrations and total RNA content quantified per gram rumen fluid (RF). Colour code indicates the four rumen-cannulated Holstein dairy cows.

### Microbiome structure and dynamics

We generated metatranscriptomes from the rumen fluid RNA using deep Illumina HiSeq paired-end sequencing and analysed the rRNA and mRNA content (Fig. 1, Supplementary Table S3). By taxonomic classification of the small subunit (SSU) rRNA transcripts we investigated if the rumen process dynamics (i.e. gas emissions and VFA production) were reflected in the taxonomic composition of the microbiome. This primer- and PCR-independent approach enables the holistic detection and classification of Eukarya, Bacteria and Archaea, typically not possible via PCR/amplicon-based techniques [28, 29]. The obtained three-domain profiles revealed that all major taxonomic groups known from ruminants were present (Fig. 2a-c, Supplementary Table S4), with eukaryotic, bacterial and archaeal taxa accounting for 25.1 ± 10.5 %, 74.5 ± 10.5 % and 0.3 ± 0.1 % of the SSU rRNA transcripts, respectively. Among the eukaryotes, Ciliates were dominant, accounting for > 70 % of SSU rRNAs in 10 out of 16 metatranscriptomes. The presence of *Entodinium* spp., *Epidinium* spp. and *Eudiplodinium maggii* and the absence of *Polyplastron multivesiculatum* was indicative of a type B ciliate community as typically found in cattle [30]. Altogether, 155 different bacterial families were detected. Out of these, 32 families were detected in all metatranscriptomes, with the ten most abundant being *Prevotellaceae, Succinivibrionaceae, Lachnospiraceae, Ruminococcaceae, Fibrobacteraceae, Spirochaetaceae, Erysipelotrichaceae, Veillonellaceae (Negativicutes),* RF16 and *Rikenellaceae,* accounting on average for 93.6 % of all bacterial SSU rRNA reads assigned to family level (Fig. 2c), potentially representing the bovine core microbiome [13]. All archaeal transcripts belonged to methanogens, with *Methanomassiliicoccales* and *Methanobacteriales* being the dominant orders.

**Figure 2.**
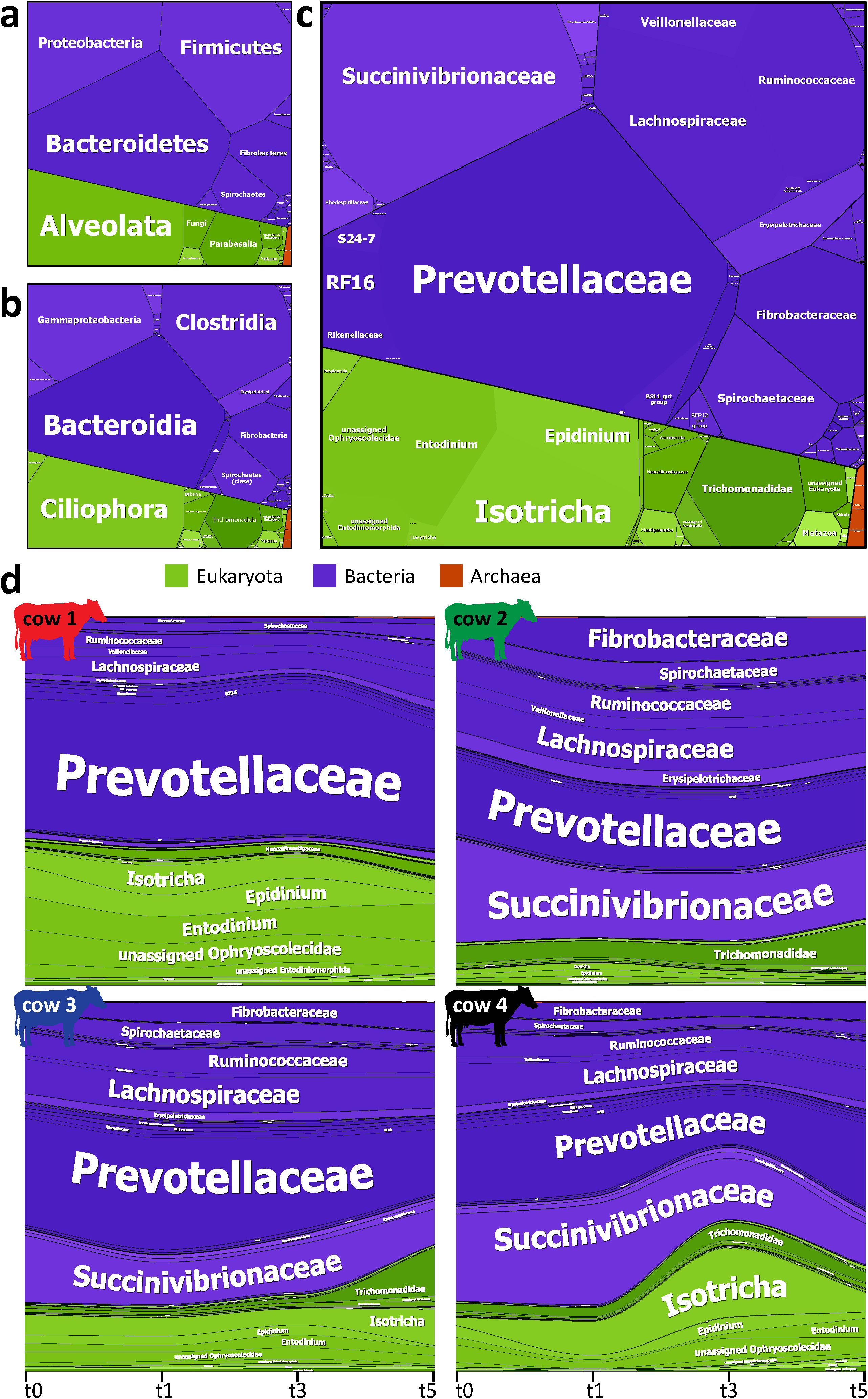
Rumen microbiome community composition and temporal dynamics. Three domain profiles showing the overall rumen microbial community composition on phylum (a), class (b) and family (c) level. Tile sizes are reflecting the average relative abundance of eukaryotic (green), bacterial (blue) and archaeal (orange) taxa observed in the 16 rumen metatranscriptomes. (d) shows the highly individual microbial communities within each cow over time. Taxa which could not be assigned on family level and/or showed relative abundance ≤ 0.01 % level are shown on higher taxonomic levels. All taxa detected in the rumen microbiomes and their relative abundances are listed in Supplementary Table S4.

Although the same major eukaryotic, bacterial and archaeal taxa were present the rumen fluid microbiomes were highly individual in each cow (Fig. 2d, Supplementary Table S4). For example, the proportion of eukaryotes in the individual rumen fluids varied from 11.6 - 40.9 % of total SSU rRNA transcripts. The two most prominent bacterial families, the *Prevotellaceae* and the *Succinivibrionaceae,* ranged from 21.0 - 66.0 % and 1.6 - 34.4 % of the microbial community composition, respectively. In contrast, their taxonomic composition was remarkably stable over the time course of the experiment (Fig. 2d) and did not show any consistent shifts in the individual cows, as revealed by several methods. Differential gene expression analysis showed no prokaryotic and no eukaryotic SSU rRNAs as differentially expressed at any time, except for three eukaryotic SSU rRNAs (i.e. *Epidinium, Eudiplodinium* and unassigned *Litostomata).* The latter were significantly less abundant at t5 compared to t3. Indicator species analysis identified several bacterial and eukaryotic taxa as significantly more abundant at certain time points (Supplementary Figure S3). However, except for the *Trichomonadidae (Parabasalia),* a group of flagellated *Protozoa,* which were found to be significantly more abundant at t5, only low abundance eukaryotic taxa were found to be indicators of the later time points. Furthermore, cow identity explained 64% of the variation in community composition (PERMANOVA p = 0.001), while time did not explain a significant amount of variation (PERMANOVA p = 0.06). Therefore, the strong temporal dynamics in rumen processes could not be explained by a successional shift in the community composition but rather by the strong increase in biomass (as reflected by the RNA content, Fig. 1d).

Analysis of mRNA gene expression profiles corroborated the notion that the observed process dynamics were an effect of overall increase in activities rather than due to an induction of specific microbial taxa or metabolisms. In two time course transitions (t3 vs. t1, t5 vs. t3), no significant differences were detected at all, while one hour after feeding less than 3 % of functional genes were significantly higher expressed (tl vs. tO). The majority (65 %) fell into the subsystem protein biosynthesis (Fig. 3, Supplementary Table S5), namely transcripts of twelve SSU and 15 large subunit ribosomal proteins and the translation elongation factor G. Additionally, relative abundance of two RNA polymerase subunit transcripts increased from tO to tl. Only very few other transcripts, involved in respiration, LPS (Kdo2-Lipid A biosynthesis) and alanine biosynthesis, biosynthesis of branched-chain amino acids, stress response, DNA repair and VFA production/consumption were significantly more abundant at tl compared to tO (Fig. 3, Supplementary Table S5). Our results suggest an immediate upregulation of the protein biosynthesis machinery as the major global response of the microbiome to feed intake.

**Figure 3.**
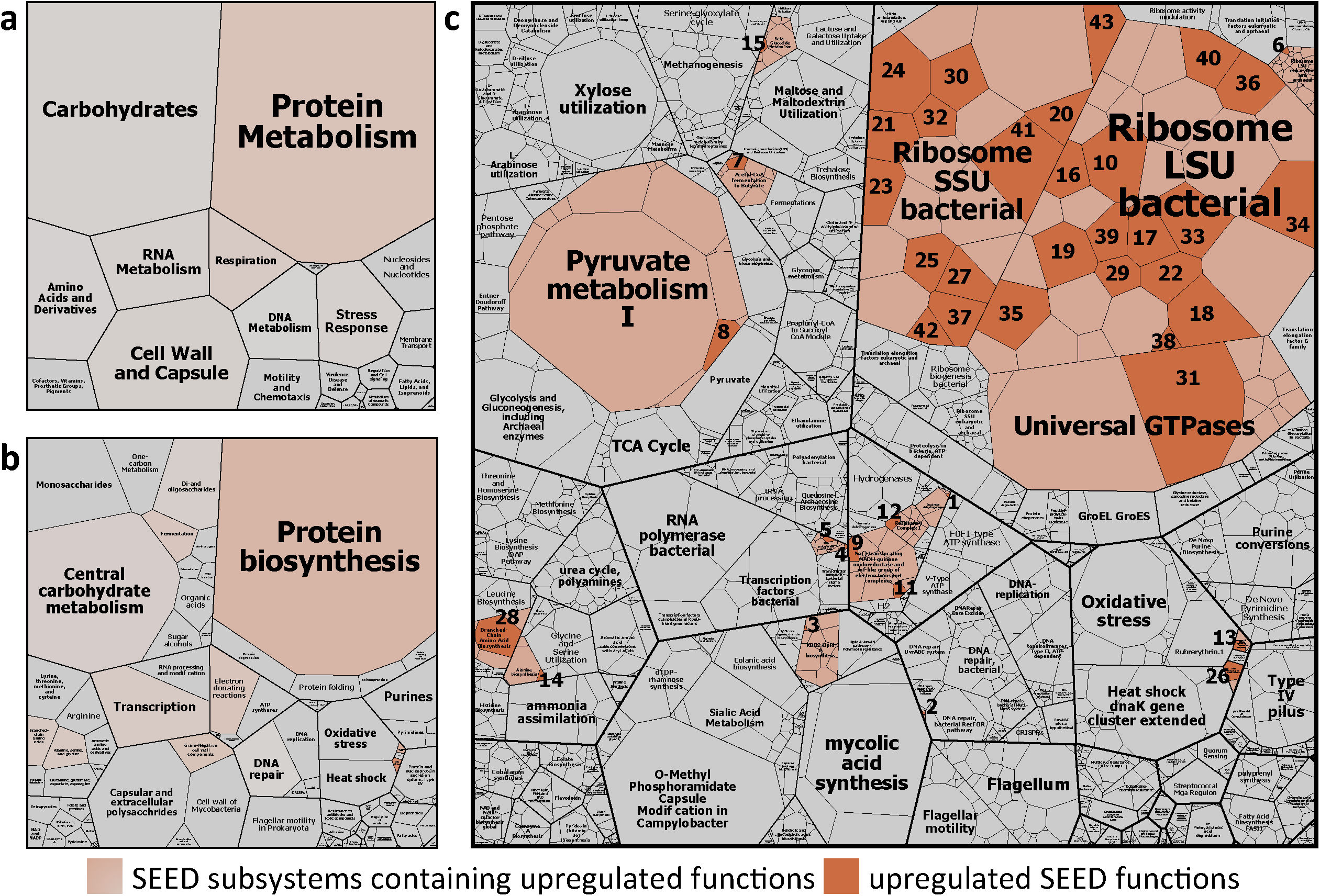
Global functional response of rumen microbiome to ruminant feed intake. Boxes showing the mean relative abundance of SEED subsystem level 1 (a), SEED subsystem level 2 (b), SEED subsystem level 3 (c) and SEED functions (c, small tiles) of eight rumen metatranscriptomes (tO and tl metatranscriptomes). Colour code indicates SEED subsystems containing functions that were identified by differential gene expression analysis to be significantly higher expressed one hour after the feeding (t1) compared to before the feeding (t0). The particular upregulated functions are coloured in orange. All functions that were subject to differential gene expression analysis (1659 SEED functions) are depicted, low abundant transcripts were excluded. For more details on the significantly higher expressed functions (e.g. functional assignment of the numbered tiles) see Supplementary Table S5.

### Quantitative metatranscriptomics

We analysed gene expression patterns of methanogens for successional changes during the experiment, which could explain the strong increase of CH_4_ emissions. However, the relative abundance of the methanogenesis-specific mRNAs and SSU rRNA transcripts of methanogen decreased at the time points with highest CH_4_ production (Fig. 4a). This pointed to a well-known problem in (meta-)omics approaches [31] i.e. linking relative abundances of taxa or genes/transcripts with biogeochemical processes that are derived from heterogeneous data. We thus calculated transcript abundances per volume of rumen fluid (equation 1) by integrating relative transcript abundance with total RNA concentrations extracted from rumen fluid. Using this quantitative metatranscriptomics approach, the transcript patterns of methanogens mirrored the observed dynamics in ruminant CH_4_ emissions, with an increase of transcripts per g rumen fluid one to three hours (t1 and t3) and a decrease five hours after the feeding started (Fig. 4b). Similar effects were observed with gene expression patterns of other, broad cellular functions, e.g. DNA replication (Supplementary Figure S4).

**Figure 4.**
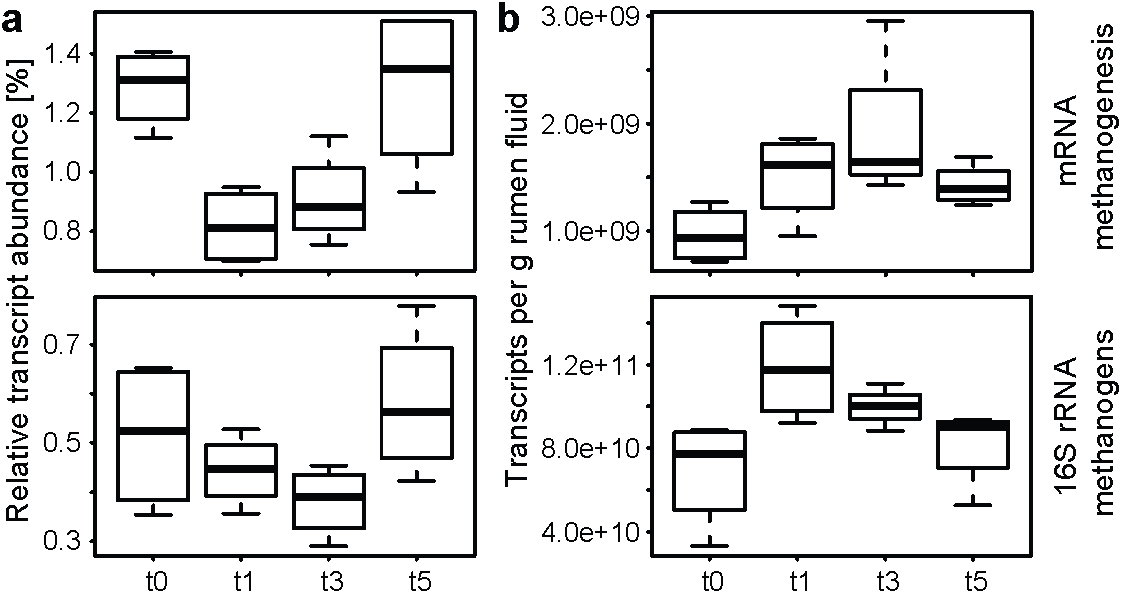
Comparison of relative and quantified transcript abundance of methanogens. Relative and quantified transcript abundance of methanogenesis specific mRNA (upper boxplots) and SSU rRNA of methanogens (lower boxplots) are depicted in (a) and (b), respectively. Data: mRNA reads assigned to the SEED subsystem Methanogenesis and SSU rRNA reads assigned to methanogens were summarized. For details on the quantification see Methods section and equation 1; x-axis: before feeding (t0), one, three, five hours after feeding started (t1, t3, t5).

### Major players in plant biomass degradation and CH_4_ production

Using this quantitative approach, we conducted a broad, integrative functional screening to identify the major microbial players in three key steps of anaerobic plant biomass degradation: (1) breakdown of complex plant polysaccharides, (2) carbohydrate fermentation to VFA and (3) methanogenesis. We used rumen fluid as proxy to analyse the complete anaerobic degradation cascade, although it has been shown that especially fibrolytic particle-associated communities can differ [32, 33].

### Degradation of plant polysaccharides

A screening for transcripts of carbohydrate active enzymes (CAZymes) revealed that the four dominant CAZyme categories were cellulases, hemicellulases, starch degrading enzymes and oligosaccharide hydrolases, accounting for 77.5 ± 2.1 % (Supplementary Figure S5 & Table S6). We quantified and taxonomically classified these transcripts to reveal their distribution among the rumen microbiome (Fig. 5). Three higher-level bacterial and two eukaryotic taxa were identified as predominantly involved, namely *Prevotellaceace (Bacteroidetes), Clostridiales (Firmicutes), Fibrobacter, Ciliophora* and *Fungi (Neocallimastigaceae).* While some of the links were known, e.g. *Fibrobacter* as major producer of cellulases [34], others are providing new insights into the complexity of CAZyme production by rumen microorganisms. For instance, Ciliates produced substantial amounts of hemicellulase and cellulase transcripts, and surprisingly few transcripts encoding starch-degrading enzymes, although they have long been considered as starch degraders [35]. Furthermore, the anaerobic fungi *Neocallimastigaceae,* produced the largest share of cellulase transcripts of all microorganisms. The abundant share of cellulase and hemicellulase transcripts encoded by *Clostridiales* establishes them as another key fibre degrading bacterial group in the rumen [36]. The data also show that *Prevotellaceae* primarily expressed genes encoding oligosaccharide hydrolases, starch degrading enzymes and hemicellulases, but not cellulases. *Firmicutes* appeared to have the broadest capacity for polysaccharide degradation, with equal abundances of CAZyme transcripts in all four investigated categories. However, the *Firmicutes (Clostridiales)* comprised several different genera within the *Ruminococcaceae* and *Lachnospiraceae,* whereas *Fibrobacteres* and *Bacteroidetes* were dominated by a single genus, *Fibrobacter* and *Prevotella,* respectively.

**Figure 5.**
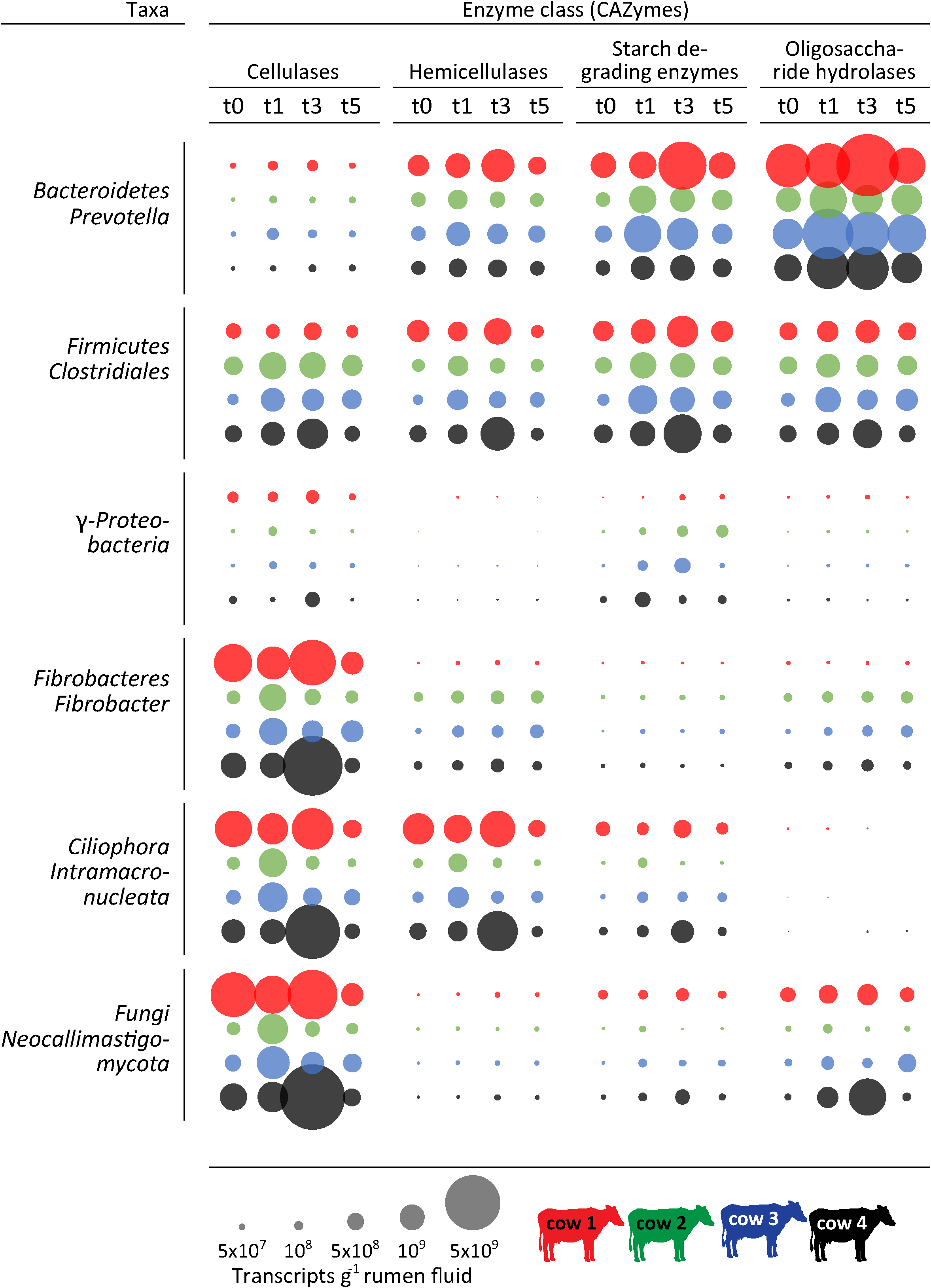
Dynamics and distribution of carbohydrate active enzymes (CAZymes) among the rumen microbiome. Circles depict the quantified numbers of CAZyme transcripts (g”^1^ rumen fluid), summarized in respective to their activity (cellulases, hemicellulases, starch degrading enzymes and oligosaccharide hydrolases), separated for the major Bacteria and Eukarya involved in the breakdown of complex plant material (on phylum level and lowest common dominant taxon). Colour code indicate the different cows and the different time points (grey scale of the columns); before feeding (tO), one, three, five hours after feeding started (tl, t3, t5).

The taxonomic distribution of CAZymes displayed strong differences between the cows, pointing to the same individuality as observed in the taxonomic composition of the rumen microbiome; e.g. Eukaryotes dominated the cellulase transcript pools in cow 1 and cow 4, whereas in cow 2 and cow 3 *Fibrobacteres* and *Firmicutes* cellulase transcripts were equally abundant to *Ciliophora* and *Fungi* cellulase transcripts. Thus, the expression of the different CAZyme categories by three to four different taxa shows a high functional redundancy for polysaccharide degradation in the rumen microbiome, within and between different domains of life.

### VFA production

Acetate, propionate and butyrate were the major VFAs accounting for 60.4 ± 4.9 %, 21.9 ± 3.4 % and 11.6 ± 2.7 % of total VFAs, respectively. VFA concentrations in the rumen fluid increased after feed intake, while the pH dropped (Fig. 1c, Supplementary Table S7). Although no VFA production or absorption rates were measured, it has been shown that VFA concentrations are suitable proxies for production rates [4]. Quantitative metatranscriptomics revealed the presence of transcripts for three complete acetate production pathways from pyruvate, i.e. directly (via pyruvate:ubiquinone oxidoreductase, *poxB),* via acetyl-CoA and via acetyl-CoA and acetyl-P (Supplementary Figure S6). The transcripts were assigned to *Bacteroidetes* (mainly *Prevotella)* and *Firmicutes* (i.e. *Clostridiales* and *Negativicutes),* with transcript levels of *Prevotella* exceeding *Firmicutes* up to 30-times in the acetyl-CoA and acetyl-P pathway (Supplementary Figure S7A and B). In general, *poxB* transcript abundances (direct conversion of pyruvate to acetate) were one to two orders of magnitude lower than abundances of the other pathways (Supplementary Figure S7C), with *Clostridiales poxB* transcripts dominating over those of *Prevotella poxB* in all samples (1.4 - 64 times). Together, these results suggest that *Prevotella* were the dominant acetate producers in this experiment.

Transcript analysis revealed the presence of two distinct pathways for propionate production (Supplementary Figure S8), (1) from succinate (succinate pathway) and (2) from lactate (acrylate pathway). Transcript levels of *Prevotella* again exceeded *Firmicutes* (i.e. *Clostridiales;* up to 20-times), suggesting that *Prevotella* also dominated propionate production (Supplementary Figure S9A). Transcripts for two complete pathways possibly leading to butyrate production were detected within the *Firmicutes,* i.e. the butyrate kinase pathway within *Clostridiales* and the butyryl-CoA:acetate CoA-transferase pathway within *Negativicutes* (Supplementary Figure S8 and S9B). These pathways only differ in the last step, i.e. the conversion of butyryl-CoA to butyrate, which is performed in two steps via butyryl-P by *Clostridiales* and directly by *Negativicutes.* In general, transcript abundances of VFA production pathway enzymes mirrored the VFA concentration patterns, especially for acetate but to a lesser extend also for propionate and butyrate (Supplementary Figure S6 & S8), with a peak in transcript abundance at t1 or t3 and a subsequent decrease of transcripts at t5. Again, the transcript abundances and their taxonomic distribution showed marked differences between the individual cows. For instance, the abundance of transcripts for acetate production via acetyl-CoA and propionate production assigned to *Bacteroidetes* (mainly *Prevotella)* was much higher in cow 1 compared to the other cows, reflecting the higher relative abundance of *Prevotella* within cow 1. Furthermore, *Negativicutes* (formerly *Veillonellaceae)* had a higher transcriptional activity for acetate production via acetyl-CoA and butyrate production than *Clostridiales* within cow 2 (Supplementary Figure S6 & S8) but not within the other cows.

### Methanogenesis

*Methanomassiliicoccales* and *Methanobacteriales* were the two dominant methanogenic orders, accounting for > 99 % of SSU rRNAs. All SSU rRNA transcripts assigned to the *Methanomassiliicoccales* belonged to the GIT clade [37] a sister lineage of *Methanomassiliicoccaceae.* Within the *Methanobacteriales,* the majority of the SSU rRNA transcripts belonged to the genus *Methanobrevibacter,* whereas *Methanosphaera* accounted for up to 13.3 % (mean 6.0 %). Between 2.7% and 24.4 % of *Methanobacteriales* SSU rRNA transcripts could not be assigned on a genus level (mean 15.2 %).

The abundance of SSU rRNA transcripts of both groups followed the CH_4_ emission dynamics (Fig. 6a). However, only *Methanomassiliicoccales* showed a strong positive linear correlation (r_s_ = 0.75, p < 0.001) and only their SSU rRNAs showed significant differences overtime similar to the CH_4_ emissions (Fig. 6a). Methyl coenzyme M reductase (Mcr), the enzyme catalysing the last step in methanogenesis is conserved in all methanogenic Archaea. The gene encoding the α-subunit of Mcr, *mcrk,* has thus been established as functional and phylogenetic marker for methanogens [38, 39]. No significant differences in *mcrk* transcript abundance were detected (Fig. 6b).

**Figure 6.**
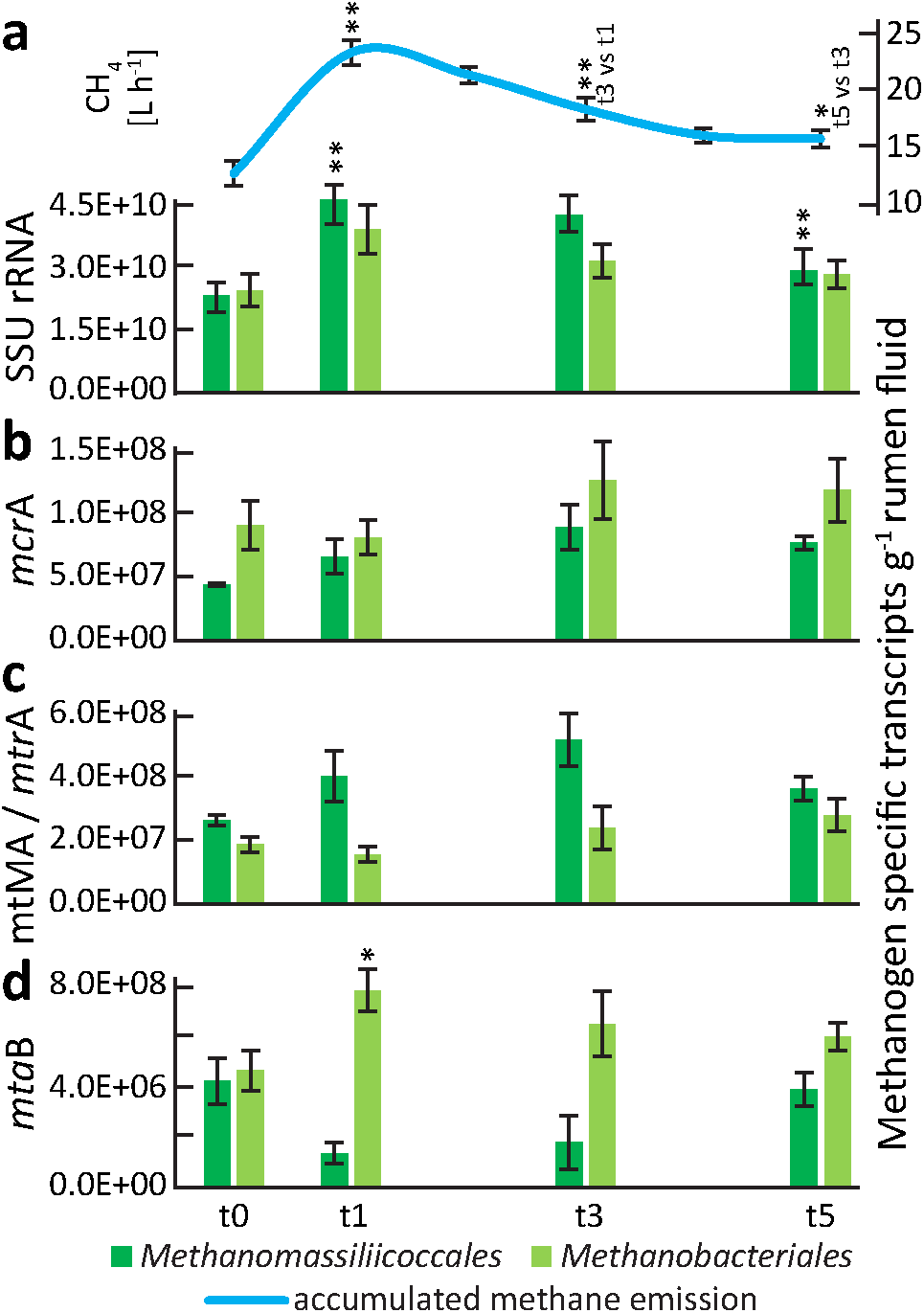
Methane and methanogen transcript dynamics during plant biomass degradation. (a) Methane emissions and quantified SSL) rRNA transcripts of the two methanogen orders present in the rumen metatranscriptomes, *Methanomassiliicoccales* and *Methanobacteriales* (i.e. *Methanobrevibacter* and *Methanosphaera),* before feeding (t0) and one (t1), three (t3) and five (t5) hours after the feeding started, (b) Quantified *mcrA* (functional marker for all methanogens) transcripts, (c) Quantified *mtMA* (methylamine-specific methyltransferases) and *mtrA* transcripts (methyl-H_4_MPT:HS-CoM methyltransferase, alpha subunit), key transcripts in *Methanomassiliicoccales* and *Methanobrevibacter* specific methanogenesis, respectively. *mtMA* summarizes mono-, di- and trimethylamine-specific methyltransferase *(mtmB, mtbB* and *mttB)* transcripts, whereas mrrB transcripts constitute to > 70 % of the *mtMA* transcripts, (d) Quantified *mtaB* (methanol-specific methyltransferase) transcripts. *Methanomassiliicoccales* and *Methanosphaera mtaB* transcripts are negatively and positively correlating with CH_4_ emissions, respectively. Mean of the four cows is shown for each time-point, error bars depict standard error of the mean (SEM). Asterisk indicate significant differences between the respective time-points and the previous one (* p < 0.5, ** p < 0.01).

The specific methanogenesis pathways differ fundamentally between *Methanomassiliicoccales* and *Methanobacteriales (Methanobrevibacter* and *Methanosphaera). Methanomassiliicoccales* are H_2_ dependent methylotrophic methanogens reducing methylamines and methanol to CH_4_with H_2_as electron donor [40, 41] In contrast, *Methanobrevibacter* produces CH_4_ mainly via the reduction of CO2 with H_2_ as electron donor. *Methanosphaera* in turn produces CH_4_ from methanol and H_2_ [42]. To identify temporal changes in the type of methanogenesis pathways actively used, we searched for transcripts of key-enzymes in these taxon-specific methanogenesis pathways: (1) Methylamine-specific methyltransferases *(mtMA),* involved in methanogenesis from methylamines by *Methanomassiliicoccales,* (2) Methyl-H_4_MPT:HS-CoM methyltransferase *(mtrA),* involved in methanogenesis from H_2_ and CO_2_ by *Methanobrevibacter* and (3) Methanol-specific methyltransferase transcripts *(mtaB)* involved in methanogenesis from methanol by *Methanosphaera* and *Methanomassiliicoccales.* We observed the same pattern for *Methanomassiliicoccales mtMA* transcripts as for the SSL) rRNA transcripts, i.e. a strong positive response to the feed intake (Fig. 6c). In contrast, no response of *Methanobrevibacter mtrA* transcript levels was observed. Immediately after feed intake, the abundance of *mtaB* transcripts of *Methanosphaera* increased, correlating positively with CH_4_ emissions (r_s_ = 0.59, p < 0.05) (Fig. 6d), while *Methanomassiliicoccales mtaB* transcripts negatively correlated with CH_4_ emissions (r_s_ = −0.63, p < 0.01). Taken together, these results indicate that only the methyl-reducing methanogens *Methanosphaera* and *Methanomassiliicoccales* responded to feed intake.

## Discussion

In this study, we used an integrated approach, combining metatranscriptomics with targeted metabolomics (gas and VFA profiling) to holistically investigate the temporal rumen microbiome dynamics during plant biomass degradation in lactating cows.

By integrating relative transcript abundances with RNA concentrations, we were able to establish the link between rumen microorganism and their activity to processes such as gas emissions and VFA production. Due to the fast growth response of the microbiome to ruminant feed intake relative transcript abundances, which are commonly used in (meta-)transcriptomics, were not sufficient to establish this link. Few studies have already applied quantitative metatranscriptomics in marine ecosystems (e.g. [43, 44]), focussing on Bacteria and nutrient cycling. Our study is the first host-associated study aiming to link process data to microbial taxa and functions. Furthermore, our approach is different as we apply total RNA concentrations instead of internal mRNA standards for sizing up metatranscriptomics. This quantitative approach allowed us to assess the temporal dynamics of major bacterial, eukaryotic and archaeal taxa involved in the three key steps of anaerobic plant biomass degradation in the cow rumen.

Our results showed that the microbiome composition was surprisingly stable during feed digestion. The strong increase of ruminant CH_4_ emissions after feeding was not related to a microbial community shift as we had hypothesized but to a fast growth response of the whole rumen microbiome. This led to enhanced fermentation rates, reflected by the increase of CO_2_, H_2_ and VFA concentrations and an associated rise in methanogenesis rates. A similar dynamic of bacterial concentrations (SSU rRNA gene copies per mL rumen fluid) as a response to ruminant feed intake was reported recently [25].

While the rumen microbiomes were stable over time, the individual microbiomes differed substantially between the four cows. Despite strong variation in abundance of bacterial and eukaryotic community members, these microbiomes exhibited similar fermentation characteristics, evidenced by gas output and VFA patterns. This points towards extensive functional redundancy among rumen microbiome members, where multiple microorganisms possess the same functional trait(s) and can replace each other [45, 13]. In fact, we could show high functional redundancy at all three key-steps of anaerobic carbohydrate degradation to CO_2_ and CH_4_.

Remarkably, inter-domain functional redundancy was widespread among the fibrolytic community, where eukaryotes and bacteria contributed varying amounts of CAZyme transcripts within individual cows. For instance, most cellulase transcripts stemmed from two bacterial *(Fibrobacter* and *Clostridiales)* and two eukaryotic groups *(Neocallimastigaceae* and *Ciliophora),* with the eukaryotes producing the largest share of cellulase transcripts in two out of the four cows. Inter-domain functional redundancy was also observed within hemicellulose, starch and oligosaccharide degradation, with marked differences between individual cows. Our results add to the growing notion that eukaryotic contribution to fibre degradation has been underestimated in the past. Very recent metatranscriptomic work with one individual sample also suggested ciliates and fungi as important for (hemi-)cellulose degradation [21].

Host individuality and functional redundancy were also revealed in the second key step of anaerobic plant biomass degradation, i.e. the fermentation of carbohydrates to VFA. Three major, well known VFA producing taxa [46, 47] were identified and their contribution to transcript pools of enzymes involved in VFA production was cow dependent. These taxa, i.e. *Bacteroidetes (Prevotella), Clostridiales* and *Negativicutes (Veillonellaceae)* produced acetate, propionate and butyrate via different fermentative pathways, of which some where shared among taxa and others were taxon-specific. Although *Prevotella* and *Clostridiales* in general dominated acetate/propionate and butyrate production, respectively, *Negativicutes* contributed substantially to acetate production via acetyl-CoA and butyrate production via the butyryl-CoA:acetate CoA-transferase pathway in cow 2.

The third and terminal step in anaerobic feed degradation is catalysed by methanogens. Also among these we observed functional redundancy. All detected groups (i.e. *Methanomassiliicoccales, Methanobrevibacter* and *Methanosphaera)* are characterised as hydrogenotrophic using H_2_as electron donor [40, 41, 42]. The removal of H_2_ is important for the rumen ecosystem and the host because low concentrations of H_2_ ensure high fermentation rates and efficient feed digestion [48]. The longitudinal experimental setup revealed temporal dynamics in electron acceptor usage within the *Methanomassiliicoccales,* where the fraction of methanol-specific methyltransferase transcripts was much lower immediately after feeding, exhibiting an opposite expression pattern to the methylamine-specific methyltransferases. In turn, it appeared that *Methanosphaera* dominated methanol reduction at these time points, showing once more the redundancy among organisms of the same functional guild. The root cause for this might be manifold, e.g. due to a higher substrate affinity of *Methanomassiliicoccales* for methylamines as compared to methanol or higher concentrations of methylamines. Alternatively, *Methanosphaera* could outcompete *Methanomassiliicoccales* for methanol under conditions of high H_2_ partial pressure. Taken together, the data suggest that the methyl-reducing *Methanomassiliicoccales* and *Methanosphaera* were responsible for the increase of CH_4_ emissions immediately after feed intake and not the CO_2_-reducing *Methanobrevibacter.* This is surprising, given that CO_2_ is a much more abundant methanogenesis substrate than methylamines and methanol. The sources of methylamines, i.e. glycine betaine (from beet) and choline (from plant membranes), and methanol (from the hydrolysis of methanolic side-groups in plant polysaccharides) are well known [49], however the amount of these substrates might vary substantially with different diets. Previous, less temporally resolved work suggested that *Methanobrevibacter* was associated with high CH_4_ emissions [14, 49]. However, a comparison of sheep rumen metagenomes and metatranscriptomes indicated that *Methanomassiliicoccales* are very active community members in both high and low CH_4_ “emitters”, with around 5 times higher abundances in the metatranscriptomes compared to the metagenomes [16]. Furthermore, their transcript abundances were significantly higher in high CH_4_ “emitters”. Also in cows, it was shown that *Methanomassiliicoccales* can represent the predominant active methanogens [24]. In fact, a need for more research on methyl-reducing methanogens in the rumen was pointed out recently [49], including quantifying their contribution to rumen methane production. Further studies on *Methanomassiliicoccales* and *Methanosphaera* physiology *in vitro* and metabolic interactions with the substrate-providing microorganisms *in situ* might identify novel targets for CH_4_ mitigation strategies, such as enzymes of the methyl-reducing pathway or the supply of methylated substrates. Such efforts might complement general methanogenesis inhibitors such as 3-nitrooxypropanol to achieve more efficient methane mitigation [50].

## Conclusions

To our knowledge, our study is the first longitudinal integrated meta-omics analysis of the rumen microbiome during plant biomass degradation. It is another step towards a comprehensive system-level understanding of the dynamic rumen ecosystem, as envisioned by Hungate and coworkers already more than 50 years ago [11]. Applying a quantitative metatranscriptomics approach, it enabled the time-resolved link between microbiome structure and function and rumen processes. It revealed a rather simple response to feed intake, namely a general growth of the whole community, without the detection of distinct successional stages during degradation. The cow-individual microbiomes exhibited a surprisingly high functional redundancy at several steps of anaerobic degradation pathway, which can be seen as example for the importance of multi-functional diversity for robustness of ecosystems, similar to what has been found in terrestrial biomes [51]. It furthermore points towards CH_4_mitigation strategies that directly tackle the producers of CH_4_, since all other functional guilds show high organismic diversity with individual taxa being replaceable by others.

## Methods

### Animal feeding trial

(Fig. 1a). The animal feeding trial was conducted at the Department of Animal Science, Aarhus University (Denmark). The animal experiments were approved by The Experimental Animal Inspectorate under The Danish Ministry of Justice (journal no. 2008/561-1500). Four rumen-cannulated lactating Holstein dairy cows were fed a typical dairy cow diet containing mainly clover grass and corn silage (Supplementary Table S1 and S2) twice a day in a semi-restrictive way. The cows were in the second parity or later, they were 215 ± 112 (mean ± standard deviation) days in milk, had live weight at 602 ± 20 kg and had a milk yield at 33.5 ± 5.4 kg (Supplementary Table S8). Prior to the sampling, which was conducted over four days, the animals had been fed the respective diet continuously for more than two weeks. Day 1: Cows were fed ad libitum. Day 2: The feed was removed at 4 am, cows were allowed to eat from 7 am to 8 am, and again from 2 pm until 4 am the next day. Rumen fluid was sampled at time points 4 am, 7 am, 8 am and every second hour until 10 pm, and with a final sampling at 4 am on day 3. Rumen fluid was randomly sampled from different areas of the rumen, pooled and filtered through sterile filter bags with a pore size of 0.5 mm (Grade Blender Bags, VWR, Denmark). The pH of the rumen liquid samples was directly analysed with a digital pH meter (Meterlab PHM 220, Radiometer, Denmark) and subsamples were frozen at −20°C for VFA analysis and other chemical analysis, or flash-frozen in liquid nitrogen and stored at −80°C for nucleic acid extraction. Day 3: Animals were transferred to custom-built transparent polycarbonate open-circuit respiration chambers (1.45 × 3.90 × 2.45 m) and fed ad libitum. Day 4: The cows were fed like on day 2. CH_4_, CO_2_ and H_2_ were quantified continuously throughout the day.

### CH_4_, CO_2_, H_2_ and VFA quantification

The open-circuit indirect calorimetry based respiration chambers, kept at slight under pressure, measured gas exchange (CH_4_, CO_2_, O_2_, and H_2_; Columbus Instruments, Columbus, USA), air flow and feed intake continuously during the experiment as described in detail in [52] and in [53]. VFAs in the rumen liquid samples were quantified using a Hewlett Packard gas chromatograph (model 6890, Agilent Technologies Inc., Wilmington, DE, USA) with a flame ionization detector and a 30-m SGE BP1 column (Scientific Instrument Services, NJ, USA) as described in [54].

### Nucleic acid extraction and linear RNA amplification

(Fig. 1a). Nucleic acids were extracted based on the method of [55], and as described in [28]. Extraction buffer and phenolxhloroform (5:1, pH 4.5, ambion), 0.5 mL of each, were added to a lysing matrix E tube (MP Biomedicals) containing approximately 0.25 g of rumen fluid sample. Cells were mechanically lysed using a FastPrep machine (MP Biomedicals, speed 5.5, 30 sec) followed by nucleic acid precipitation with PEG 8000. All steps were performed on ice or at4°C. Nucleic acids were re-suspended in 50 μL DEPC H_2_O and 1 μL of RNaseOUT^™^ (Thermo Fisher Scientific) was added. 10 μL of nucleic acid extracts were subject to DNase treatment (RQ1 DNase, Promega) and subsequent RNA purification (MEGAclear^™^ Kit, Ambion). Quantity and quality of RNA was assessed via agarose gel electrophoresis, NanoDrop^®^ (ND 1000, peqlab) and Qubit^™^ (Thermo Fisher Scientific). Absence of DNA in the RNA preparations was verified by PCR assays targeting bacterial SSU rRNA genes and archaeal *mcrA* genes. MessageAmp^™^ Il-Bacterial Kit (ambion) was used according to the manufacturer’s manual to synthesise cDNA (via polyadenylation of template RNA and reverse transcription) and perform in *vitro* transcription on the cDNA to amplify total RNA.

### Sequencing and sequence data pre-processing

Illumina HiSeq 2500 paired-end (125 bp) sequencing was performed at the Next Generation Sequencing Facility of the Vienna Biocenter Core Facilities on cDNA. The template fragment size was adjusted that paired sequence reads could be overlapped. We used PRINSEQ lite v. 0.20.4 [56] to apply quality filters and trim the reads (parameters -minjen 180 -min_qual_mean 25 -ns_max_n 5 -trim_tail_right 15 -trim_tail_left 15). SortMeRNA v. 2.0 [57] was used to separate sequence reads into SSU rRNA, LSU rRNA and putative mRNA reads. For more details and results of the initial data processing steps see supplementary Table S3. All computations were performed using the CUBE computational resources, University of Vienna (Austria), or run on the HPC resource STALLO at the University of Troms0 (Norway). Raw sequence data have been submitted to the NCBI Sequence Read Archive (SRA) under the accession numbers SAMN07313968 -SAMN07313983.

### Taxonomic classification of SSU rRNA reads

We generated random SSU rRNA subsamples containing 50,000 reads out of all SSU rRNA reads with a length between 200 to 220 bp (45.8 ± 11.5 % of total SSU rRNA reads). These subsamples were taxonomically classified with BLASTN against the SilvaMod rRNA reference database of CREST [58] and analysed with MEGAN [59] v. 5.11.3 (parameters: minimum score 100, minimum support 1, top 2 %, 50 best blast hits). Three domain profiles were visualised with treemaps based on CREST taxonomy. Statistical analyses were done with R [60]; packages: edgeR [61], vegan [62], indicspecies [63], heatmap3 [64].

### Analysis of mRNA

All putative mRNA reads were compared against the GenBank nr database using DIAMOND ([65]; v0.7.11, database as of December 2015, CUBE).

### CAZzymes

Randomly selected subsamples of 2 million nucleotide reads per dataset were translated into open reading frames (ORFs) of 30 amino acids or longer. The ORFs were screened for protein families using HMMER and reference Hidden Markov Models (HMMER v3.0, against the Pfam database v27; [66]). All database hits with e-values below a threshold of 10^M^ were counted. Pfam annotations were screened for CAZymes using Pfam models of previously identified CAZymes [67] and additional rumen relevant CAZymes [22] as well as CAZymes added to the Pfam-A database after these publications and summarized into higher categories (Supplementary Table S6). Translated reads assigned to any Pfam model of one of the four most dominant categories i.e. cellulases, hemicellulases, starch degrading enzymes and oligosaccharide hydrolases (Supplementary Figure S5) were extracted and blastp was used to obtain taxonomic information (blastp against the monthly updated nr db 04.2016, CUBE). BLAST tables were imported in MEGAN (parameters: minimum score 50, minimum support 1, top 5 *%,* 25 best blast hits) and further analysed. CAZymes were quantified as described below (equation 1).

### VFA

All mRNA reads assigned to any major taxa involved in the production of VFA, as identified by the SEED analyzer implemented in MEGAN, were subject to further analysis to reconstruct major VFA production (turnover) pathways. These metatranscriptomic libraries were screened for all enzymes (via their respective EC numbers) involved in the production/turnover of acetate, propionate and butyrate, by blastp searches (evalue threshold le”^10^) using the metatranscriptomic libraries as queries against the UniRef50 database (montly updated, 12.2016, CUBE). The respective enzyme names were derived from the KEGG reference pathways and literature [68, 69]. Heatmaps were constructed in R using quantified data (ng transcripts g” rumen fluid; equation 1) normalized by the sum of each transcripts over all time-points for each individual cow.

### Methanogenesis

Specific transcripts for methanogenesis were extracted from the DIAMOND annotation files via MEGAN and the implemented SEED analyzer. Assignments were critically manually evaluated and in case of uncertainty blastn was used to verify accuracy and origin of the methanogenesis transcripts as well as of the SSU rRNA transcripts (against the NCBI and Silva databases as of September 2016). Transcripts were quantified (equation 1). Pearson’s product-moment correlations and spearman rank correlation coefficients (rho = r_s_) between methanogen specific transcripts (pathway specific key transcripts and SSU rRNA transcripts) were calculated and paired t-test was used to assess temporal differences in transcript abundance (R functions: shapiro.test, contest, t.test).

### Differential gene expression analysis

mRNA. DIAMOND annotations were imported in MEGAN (parameters: minimum score 40, minimum support 1, top 10 %) and relative abundances of mRNA reads assigned to a SEED function were subject to differential gene expression analysis using edgeR (function: glmFit). Low expressed genes were filtered out and the default TMM method was used to normalize the data. To account for the cow differences a design matrix was constructed prior to the analysis to account for our experimental design and correct for batch effects (cow differences). rRNA. Taxon tables, as described above were subject to differential gene expression analysis following the same workflow as described for the SEED functions.

### Quantification of mRNA and rRNA transcripts per gram rumen fluid

We quantified mRNA and rRNA transcripts per gram rumen fluid as follows:

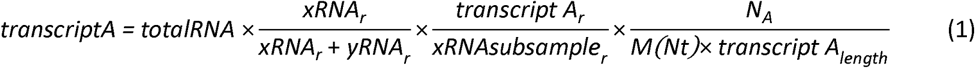

where *totalRNA* is the amount of RNA [μg] extracted per gram rumen fluid, *xRNA_r_, yRNA_r_* and *xRNAsubsample_r_are* the number of reads of m/rRNA, r/mRNA and m/rRNA subsample used for functional annotation or taxonomic classification, respectively. *transcriptA_r_* and *transcriptAi_ength_* are the number of reads assigned to a certain transcript and the length of the particular transcript. *N_A_* is the Avogadro constant and *M(Nt)* is the average molecular weight of a ssDNA nucleotide (330 × 10^6^ ug mol”^1^). For the transcript lengths we used average values of 1000 and 1500/1900 (prokaryotes/eukaryotes) nucleotides for mRNA and rRNA transcripts, respectively. As previously observed [70] the polyadenylation during cDNA synthesis is moderately enriching mRNA, therefore a ratio of mRNA:totalRNA reads of 1:25 was used to calculate transcript numbers per gram rumen fluid, as this ratio was observed in a previous study on the rumen microbiome of cows from the same breed, housed at the same facility, fed a diet containing similar amounts of neutral detergent fibre, crude protein and fat [24].

## Declarations

### Ethics approval and consent to participate

The animal experiments were approved by The Experimental Animal Inspectorate under The Danish Ministry of Justice (journal no. 2008/561-1500).

### Consent for publication

Not applicable.

### Availability of data and material

Raw sequence data have been submitted to the NCBI Sequence Read Archive (SRA) under the accession numbers SAMN07313968 - SAMN07313983.

### Competing interests

The authors declare that they have no competing interests.

### Funding

A.S. was financially supported by a scholarship of the University of Vienna for doctoral candidates (uni:docs), a travel scholarship of the University of Vienna (KWA) and a scholarship by the OeAD (Austrian agency for international mobility and cooperation in education, science and research) for doctoral candidates (Marietta-Blau-Fellowship). T.U acknowledges financial support from the University of Greifswald.

## Authors’ contributions

The study was designed by M.P., A.S. and T.U. Samples were collected by M.P. Gas and VFA quantifications were established and performed by M.P. AL.H. and P.L. RNA extractions and sample preparation were performed by A.S. Analysis of Illumina sequencing data were performed by A.S., AT.T. and T.U. Statistical analyses and figures were done by A.S. and J.B, assisted by M.B. and AT.T. The manuscript was written by A.S. and T.U., assisted by all co-authors.

## Acknowledgements

We thank Thomas Rattei (University of Vienna) for bioinformatics support and Andreas Sommer and the Vienna Biocenter Next Generation Sequencing Facility (www.vbcf.ac.at) for library preparation and sequencing.

## Supplementary Information

Supplementary Information is provided in the PDF document “Supplementary_Figures_Tables.pdf”. This Supplementary Information contains Figures and Tables supplementing the main manuscript.

